# Pharmacological GCase Activity Enhancement Inhibits Tau Accumulation

**DOI:** 10.1101/2024.08.13.607706

**Authors:** Matteo Ciccaldo, Natàlia Pérez-Carmona, Ester Piovesana, Sara Cano-Crespo, Ana Ruano, Aida Delgado, Ilaria Fregno, Beatriz Calvo-Flores Guzmán, Manolo Bellotto, Maurizio Molinari, Joanne Taylor, Stéphanie Papin, Ana María García-Collazo, Paolo Paganetti

**Affiliations:** Laboratory for Aging Disorders, Laboratories for Translational Research, Ente Ospedaliero Cantonale, Bellinzona, Switzerland; Gain Therapeutics Sucursal en España, Parc Científic de Barcelona, Barcelona, Spain; Institute for Research in Biomedicine, Faculty of Biomedical Sciences, Università della Svizzera italiana, Bellinzona, Switzerland; GT Gain Therapeutics SA, Lugano, Switzerland; School of Life Sciences, École Polytechnique Fédérale de Lausanne, Lausanne, Switzerland; Gain Therapeutics Inc., Bethesda, USA; Faculty of Biomedical Sciences, Università della Svizzera Italiana, Lugano, Switzerland

## Abstract

A slow decline in the autophagy-lysosomal pathway is a hallmark of the normal aging brain. Yet, an acceleration of this cellular function may propel neurodegenerative events. In fact, mutations in genes associated with the autophagy-lysosomal pathway can lead to Parkinson’s disease. Also, amyloidogenic protein deposition is observed in lysosomal storage disorders, which are caused by genetic mutations representing risk factors for Parkinson’s disease. For example, Gaucher’s disease *GBA1* mutations leading to defects in lysosomal sphingolipid metabolism cause α-synuclein accumulation. We observed that increased lysosomal Tau accumulation is found in human dermal fibroblasts engineered for inducible Tau expression. Inhibition of the *GBA1* product GCase augmented Tau-dependent lysosomal stress and Tau accumulation. Here, we show increased Tau seed-induced Tau accumulation in Gaucher’s fibroblasts carrying *GBA1* mutations when compared to normal fibroblasts. Pharmacological enhancement of GCase reversed this effect, notably, also in normal fibroblasts. This suggests that boosting GCase activity may represent a therapeutic strategy to slow down aging-dependent lysosomal deficits and brain protein deposition.

## Introduction

Neurodegenerative tauopathies are characterized by the progressive accumulation of neurofibrillary hyperphosphorylated Tau protein in the brain^1–3^. Tau pathology can occur secondarily to the deposition of other amyloidogenic proteins forming e.g., β-amyloid plaques in Alzheimer’s disease or α-synuclein Lewy bodies in Parkinson’s disease^4^. Protein deposition is linked to a deterioration of cellular functions eventually leading to cell death. A slow decline in cell functions is a hallmark of normal aging as observed e.g., for the autophagy-lysosomal pathway (ALP). This important cellular pathway is dedicated to the elimination of intracellular organelles and macromolecules as well as protein aggregates^5,6^. Not surprisingly, ALP malfunction contributes to the neurodegenerative process. Indeed, the deposition of amyloidogenic proteins has been reported in lysosomal storage disorders (LSD)^7,8^. LSD are caused by inherited defects in lysosomal or non-lysosomal proteins resulting in aberrant buildup of lysosomal substrates and deleterious ALP function^9^. The correlation between protein accumulation and ALP malfunction is better documented in Parkinson’s disease. Whereas monoallelic mutations in ALP genes may represent risk factors for Parkinson’s disease, homozygous mutations cause LSD^10,11^. An important example is given by the *GBA1* gene encoding for glucocerebrosidase (GCase), a lysosomal enzyme metabolizing glucosylceramide. In fact, biallelic *GBA1* mutation causes Gaucher’s disease, a mendelian LSD disorder affecting several organs and tissues due to cells accumulating fatty substances. Yet, monoallelic *GBA1* variants stand for the main genetic risk for Parkinson’s disease, suggesting that GCase malfunction may be linked to protein accumulation in the brain^12^. Loss of GCase activity in *GBA1*-Parkinson’s patients has been observed in brain, blood, and cells such as dermal fibroblasts or dopaminergic neurons derived from pluripotent stem cells^13–15^. Indicating a driving role of ALP malfunction – and possibly also of aging – in the pathogenesis of Parkinson’s disease, *GBA1* mutation carriers show defects in sphingolipid metabolism and α-synuclein accumulation^16,17^. Noteworthy, lysosomal GCase malfunction appears to occur also in the idiopathic Parkinson brain^18^ and fibroblasts^19^. Decreased GCase and buildup of glucosylceramide in degradative organelles (DOs) can inhibit autophagy^20^ contributing to α-synuclein accumulation, which in turn disturbs the targeting of GCase to lysosomes^21^ and interferes directly with GCase activity^22,23^.

We recently showed that this noxious circle of events also occurs in relation to Tau accumulation^24^. When treated with extracellular Alzheimer’s brain-derived Tau seeds, a dramatic increase in Tau accumulation in DOs was observed in primary human dermal fibroblasts engineered for inducible fluorescently tagged Tau expression. In this cellular system, Tau was shown to be a substrate of macroautophagy and to accumulate in DOs causing lysosomal stress monitored by nuclear TFE3 translocation and accumulation of lactosylceramide, a product of glucosylceramide metabolism. Notably, pharmacological GCase inhibition augmented Tau-dependent lysosomal stress and Tau accumulation^24^. The use of primary dermal fibroblasts allows for exploring whether the *GBA1* genotype may affect Tau-dependent pathological phenotypes. So, in the current study, we show increased seed-induced Tau accumulation in Gaucher’s fibroblasts carrying the *GBA1^L444P/L444P^* biallelic mutation when compared to *GBA1^WT^* fibroblasts. Gain Therapeutics has applied its drug discovery platform to develop pharmacological chaperones that allosterically bind GCase and enhance its activity, such as GT-02216^25,26^. Tau accumulation was rescued by GT-02216 on the *GBA1^L444P/L444P^* and on the *GBA1^WT^* genotype.

## Results

### GT-02216 binds to the GCase protein

Gain Therapeutics has applied its proprietary SEE-Tx® drug discovery platform ^25,27,28^ to identify a druggable allosteric site of GCase distinct from the active pocket and for performing virtual compound screening. This methodology starts from the high-resolution structure of native human GCase (PDB code: 2V3F^29^) to find novel sites and binding hotspots for guiding molecular docking^30^. The virtual screening was performed with a non-redundant library of 4.8 million commercial drug-like compounds. The best scoring compounds were experimentally validated and served as a starting point for a medicinal chemistry program that led to the discovery of GT-02216^26^ (**Fig. 1A**).

**Figure 1.**
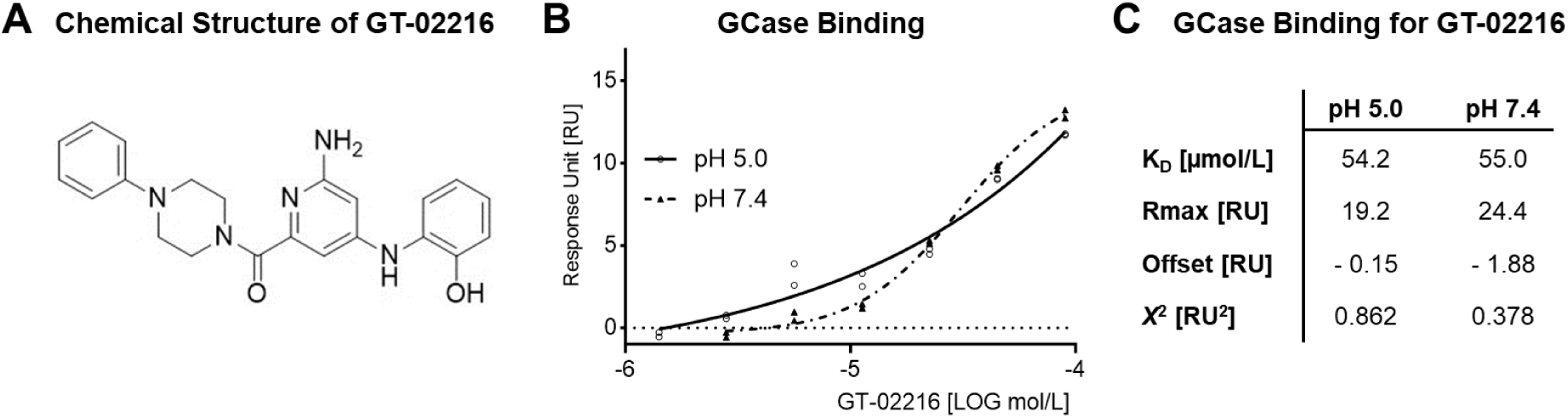
**A.** Molecular structure of the GCase pharmacological chaperone GT-02216. **B.** SPR dose-response for GT-02216 binding to immobilized human GCase protein monitored at acidic (pH 5.0) and neutral (pH 7.4) conditions. **C.** SPR binding properties determined at the indicated pH values.

The direct binding of GT-02216 to GCase was confirmed by surface plasmon resonance (SPR). SPR allows the study of the strength and kinetics of molecular interactions in real time^31^. Performing SPR experiments at pH 5.0 and pH 7.4, can provide insight into how molecular interactions behave under different physiological conditions, mimicking e.g., acidic organelles or the endoplasmic reticulum, respectively^32^. GT-02216 binds to GCase protein in a dose-dependent manner at both acidic pH 5.0 and neutral pH 7.4 (**Fig. 1B**) with similar binding affinity (**Fig. 1C**).

### GT-02216 enhances GCase activity in primary human fibroblasts

The presence of *GBA1* mutations has a deleterious effect on GCase activity. This was first confirmed in fibroblasts using an enzyme activity assay. In this assay, the fluorescent product 4-methylumbelliferone (4MU) is released by GCase activity from the synthetic substrate 4-methylumbelliferyl-β-D-glucopyranoside. Basal GCase activity was found to be reduced for all mutant *GBA1* fibroblasts analyzed in a panel of dermal fibroblasts derived from Gaucher’s patients (**Fig. 2A**).

**Figure 2.**
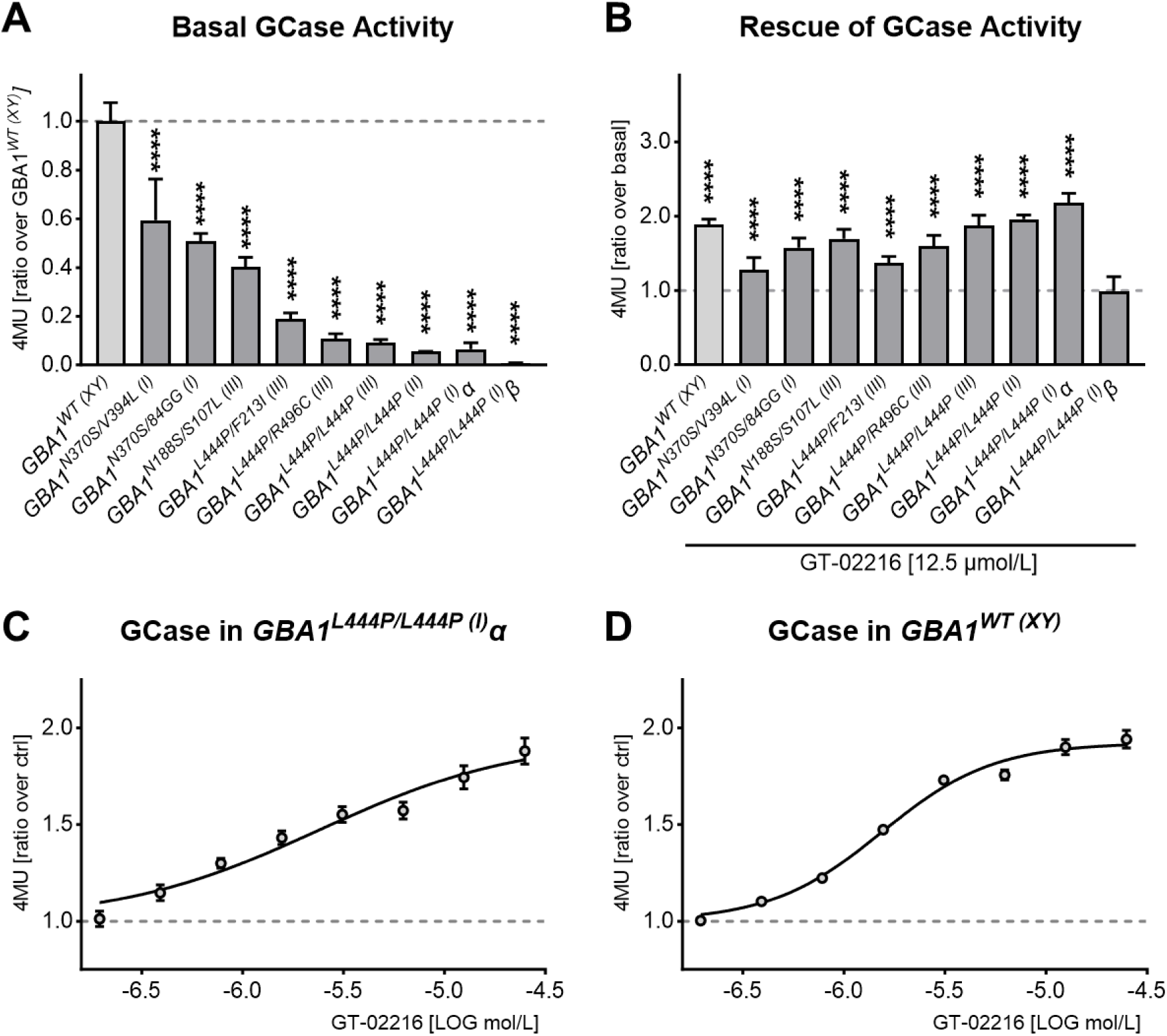
**A.** Basal GCase activity in wild-type and mutant GBA1 fibroblasts (mean ± SD normalized on GBA1^WT(XY)^, n= 6-48). Ordinary 1way ANOVA (p < .0001) and Dunnet’s multiple tests against GBA1^WT(XY)^, **** p < .0001. **B.** Effect of 4-days treatment with 12.5 µmol/L GT-02216 on GCase activity on fibroblast lines (mean ± SD normalized on the respective basal activity, n= 5-42). Ordinary 1way ANOVA (p < .0001) and Šidák’s multiple tests against the respective basal activity, **** p < .0001. **C.** GT-02216 dose-response on GBA1^L444P/L444P(I)^α fibroblasts (mean ± sem normalized on vehicle, n= 14) or **D.** on GBA1^WT (XY)^ fibroblasts (mean ± sem normalized on vehicle, n= 8) treated for 4 days. Non-linear fit with four parameters log(agonist); EC_50_= 2.4 μmol/L on GBA1^L444P/L444P(I)^α (1.3-10.5 μmol/L 95% confidentiality interval, top 1.8-2.4 μmol/L, R^2^= 0.71) and 1.5 μmol/L on GBA1^WT(XY)^ (1.3-1.8 μmol/L 95% confidentiality interval, top 1.9-2.0 μmol/L, R^2^= 0.95).

Treatment of the cells with the pharmacological chaperone GT-02216 for 4 days resulted in a significant enhancement of GCase activity in mutant *GBA1* fibroblasts as well as in *GBA1^WT(XY)^* fibroblasts (**Fig. 2B**), except for the *GBA1^L444P/L444P^*^(I)^β cell line which showed undetectable GCase activity (**Fig. 2A** and **B**). GT-02216 elicited maximal, ∼2-fold increased activity on the fibroblast lines harboring the homozygote *GBA1^L444P/L444P^* mutation. Consequently, the *GBA1^L444P/L444P^*^(I)^α line and a *GBA1^WT^*^(XX)^ line were chosen as the cellular models for subsequent studies. Treatment of these two fibroblast lines with increasing concentrations of GT-02216 for 4 days resulted in a dose-dependent rescue of GCase activity (**Fig. 2C** and **D**).

Reduced GCase activity is expected to induce an accumulation of the primary physiological GCase substrates glucosylceramide (GlcCer) and glucosylsphingosine. In fibroblasts, GlcCer is usually evaluated through the measure of hexosylceramide (HexCer), a racemic mixture composed of GlcCer and galactosylceramide. Indeed, treatment of *GBA1*^L^*^444P/L444P^*^(I)^α fibroblasts with conduritol B epoxide (CBE), a well-known potent irreversible inhibitor of GCase^33^, led to a massive accumulation of HexCer as previously described in the literature^34^ (**Fig. 3A**). Notably, GT-02216 reduced HexCer in the same mutant fibroblast line (**Fig. 3B**). In line with these data, the amount of HexCer measured in *GBA1^WT^*^(XX)^ fibroblasts was much lower compared to that present in *GBA1^L444P/L444P^*^(I)^α fibroblasts (**Fig. 3C**). GT-02216 treatment decreased HexCer also in the wild-type fibroblasts (**Fig. 3D**), consistent with the enhancement of wild-type GCase measured in the 4MU activity assay (**Fig. 2A** and **D**).

**Figure 3.**
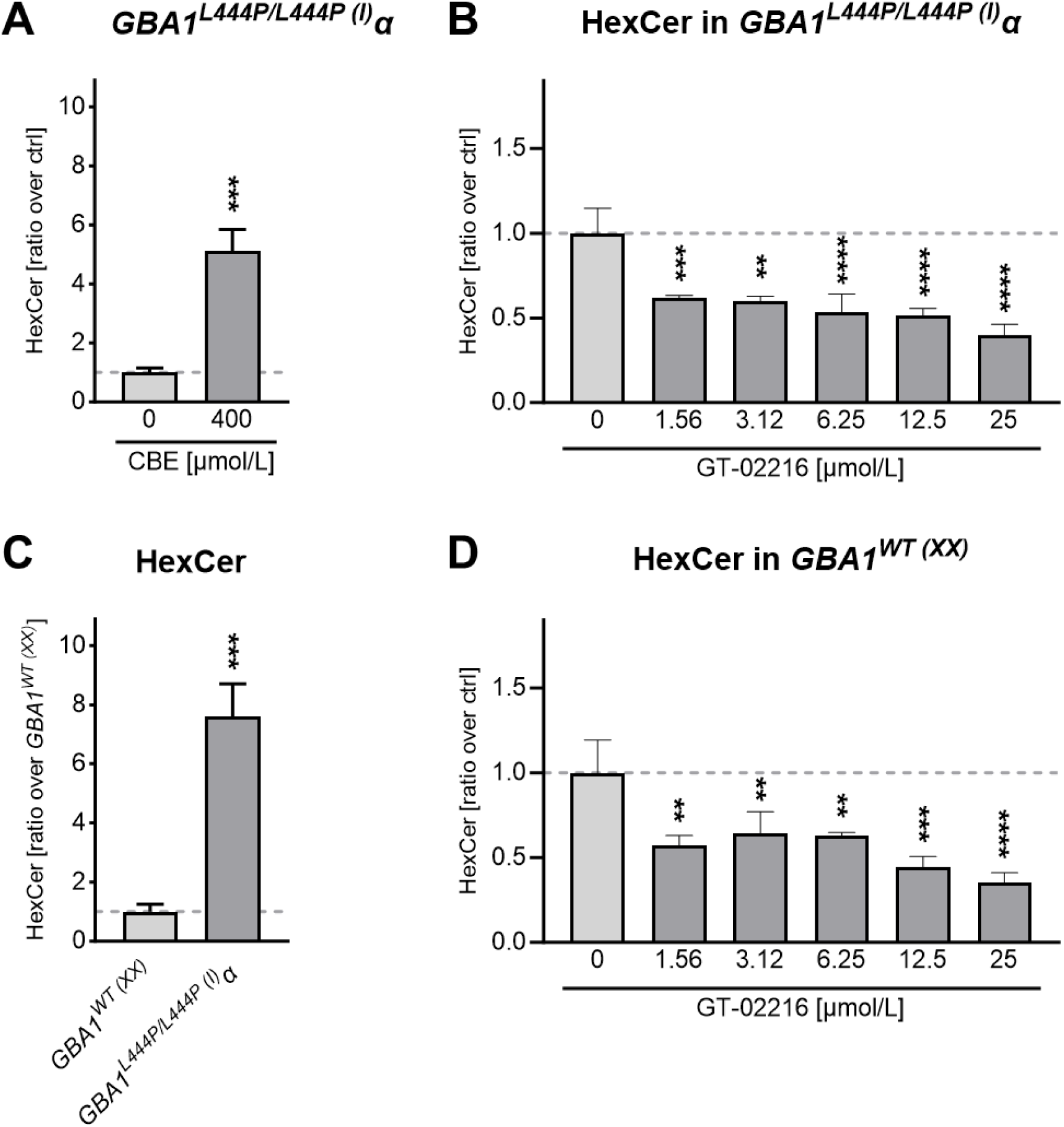
**A.** Quantification of total HexCer in GBA1^L444P/L444P(I)^α fibroblasts treated with vehicle or CBE (mean ± SD normalized on ctrl, n= 3). Unpaired t test, *** p .0007. **B.** GBA1^L444P/L444P(I)^α fibroblasts treated for 10 days with the indicated amount of GT-02216 (mean ± SD normalized on vehicle, n=3). Ordinary 1way ANOVA (p < .0001) and Dunnett’s multiple tests against vehicle. ** p < .01, *** p < .001, **** p < .0001. **C.** Quantification of total HexCer in GBA1^L444P/L444P(I)^α compared to GBA1^WT(XX)^ fibroblasts (mean ± SD normalized on GBA1^WT(XX)^, n= 3). Unpaired t test, p .0006. **D.** As in **B.** for GBA1^WT(XX)^ fibroblasts. Ordinary 1way ANOVA (p .0002) and Dunnett’s multiple tests against vehicle. ** p < .01, *** p < .001, **** p < .0001.

Next, we confirmed the data obtained for the parental human dermal fibroblasts in the two lines, *GBA1^L444P/L444P^*^(I)^α and *GBA1^WT^*^(XX)^, engineered for inducible Tau-mCherry expression. Upon induction of ectopic Tau expression with doxycycline, Tau-*GBA1^L444P/L444P^*^(I)^α fibroblasts presented a 17.5% of the GCase activity measured in Tau-*GBA1^WT^*^(XX)^ fibroblasts (**Fig. 4A**). We confirmed the ability of the pharmacologic chaperone GT-02216 to rescue GCase activity in a dose-dependent manner in Tau-*GBA1^L444P/L444P^*^(I)^α fibroblasts (**Fig. 4B**). GT-02216 boosted GCase activity in Tau-*GBA1^WT^*^(XX)^ fibroblasts independently of the absence or presence of Tau expression (**Fig. 4C**).GT-02216 also elicited a dose-dependent increase in GCase activity also in Tau-*GBA1^WT^*^(XX)^ fibroblasts (**Fig. 4D**). The dose-response curves differed somehow from those obtained in parental fibroblasts (**Fig. 2C** and **D**), and one explanation could be that the experiments were conducted in different laboratories.

**Figure 4.**
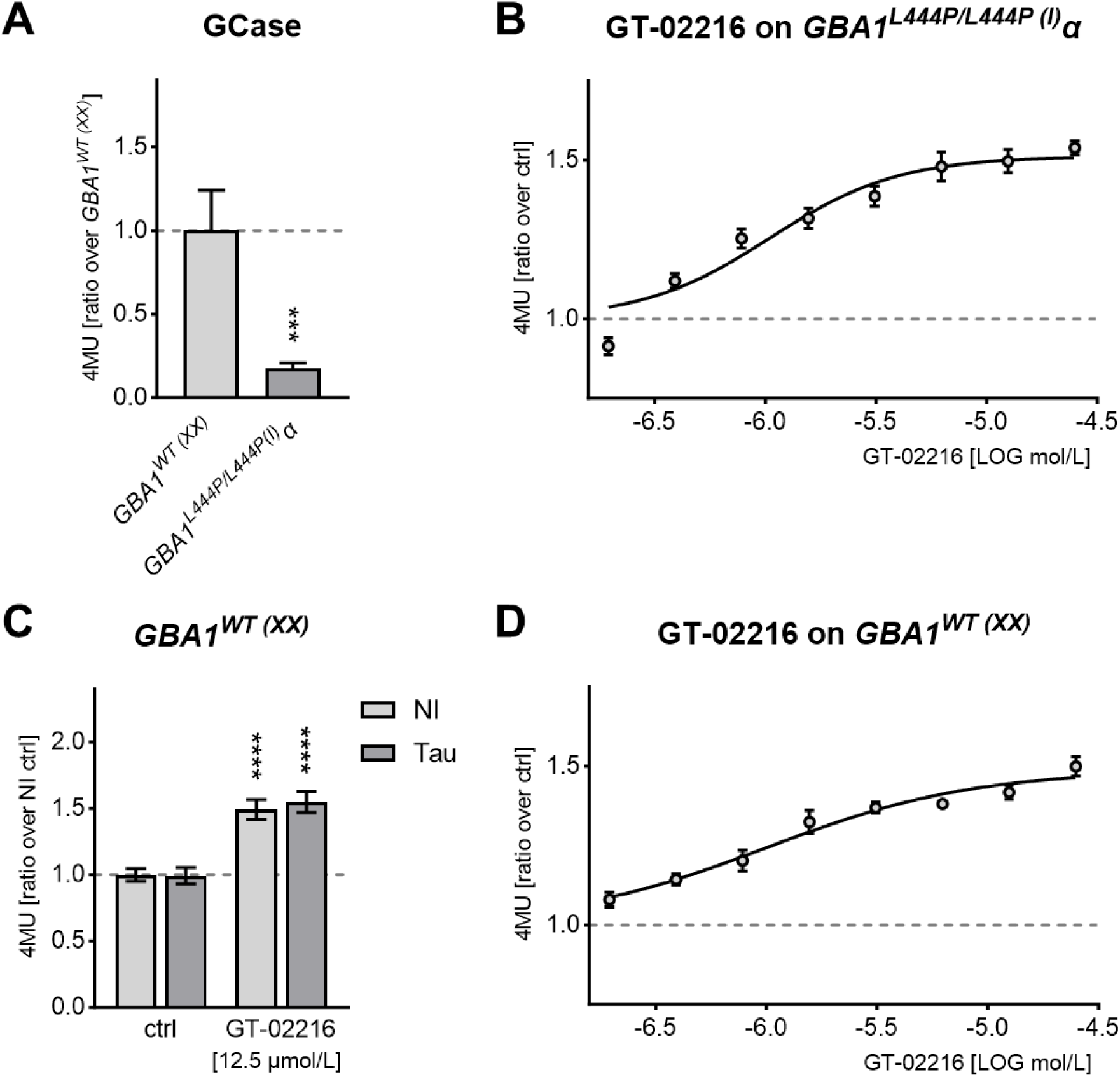
**A.** GCase activity in doxycycline-inducible Tau-mCherry human fibroblasts with the genotype GBA^WT(XX)^ or GBA1^L444P/L444P(I)^α (mean ± SD normalized on GBA^WT(XX)^, n= 7). Mann-Whitney test, *** p .0006. **B.** GT-02216 dose-response on Tau-GBA1^L444P/L444P(I)^α fibroblasts (mean ± sem normalized on vehicle, n= 8) treated for 4 days. Non-linear fit with four parameters log(agonist); EC_50_= 1.1 μmol/L (0.7-1.6 μmol/L 95% confidentiality interval, top 1.5-1.6 μmol/L, R^2^= 0.79). **C.** Basal and GT-02166-rescued GCase activity is not affected by the induction of Tau-mCherry expression with doxycycline in Tau-GBA1^WT(XX)^ fibroblasts (mean ± SD normalized on basal ctrl, n= 3). 2way ANOVA (p <.0001 for treatment, ns for Tau expression) and Šidák’s multiple tests against the respective ctrl, **** p < .0001. **D.** GT-02216 dose-response on Tau-GBA1^WT(XX)^ fibroblasts (mean ± sem normalized on vehicle, n= 8) treated for 4 days. Non-linear fit with four parameters log(agonist); EC_50_= 1.0 μmol/L (0.7-2.3 μmol/L 95% confidentiality interval, top 1.4-1.6 μmol/L, R^2^= 0.78).

### Increased Tau accumulation in mutant *GBA1* fibroblasts is reduced by GT-02216

We reported previously Tau accumulation in DOs of human primary fibroblasts treated with extracellular brain-derived Tau seeds. Moreover, pharmacological GCase inhibition with CBE increased Tau accumulation and induced lysosomal stress, suggesting a synergistic effect between lysosomal stress and the presence of accumulated Tau in DOs^24^. So, next we asked whether the presence of mutant GCase might affect Tau accumulation in DOs of primary human fibroblasts. Tau accumulation at basal conditions was 3.5-fold higher in Tau-*GBA1^L444P/L444P^* ^(I)^α fibroblasts compared to Tau-*GBA1^WT^*^(XX)^ fibroblasts (**Fig. 5A**). Moreover, when Tau-*GBA1^L444P/L444P^*^(I)^α fibroblasts were treated with Alzheimer brain-derived Tau seeds, Tau accumulation increased by a further 2.7-fold (**Fig. 5B**), but the same treatment did not affect GCase activity (**Fig. 5C**). Tau-*GBA1^WT^*^(XX)^ fibroblasts displayed a lower, ∼1.6-fold seed-induced Tau accumulation when compared to basal conditions (**Fig. 5D**), in the absence of altered GCase activity (**Fig. 5E**). These data show that genetic GCase impairment potentiated seed-induced Tau accumulation. However, treatment of Tau-*GBA1^L444P/L444P^*^(I)^α fibroblasts for 4 days with GT-02216 resulted in a dose-dependent reduction of Tau accumulation both at basal (no seeds) or in the presence of Tau seeds (**Fig. 5F**). GT-02216 reduced Tau accumulation also in Tau-*GBA1^WT^*^(XX)^ fibroblasts although the dose-dependency showed reduced potency (**Fig. 5G**). Overall, the data indicate that intracellular Tau accumulation was dependent on lysosomal function that was affected by the enzymatic activity of a single lysosomal enzyme.

**Figure 5.**
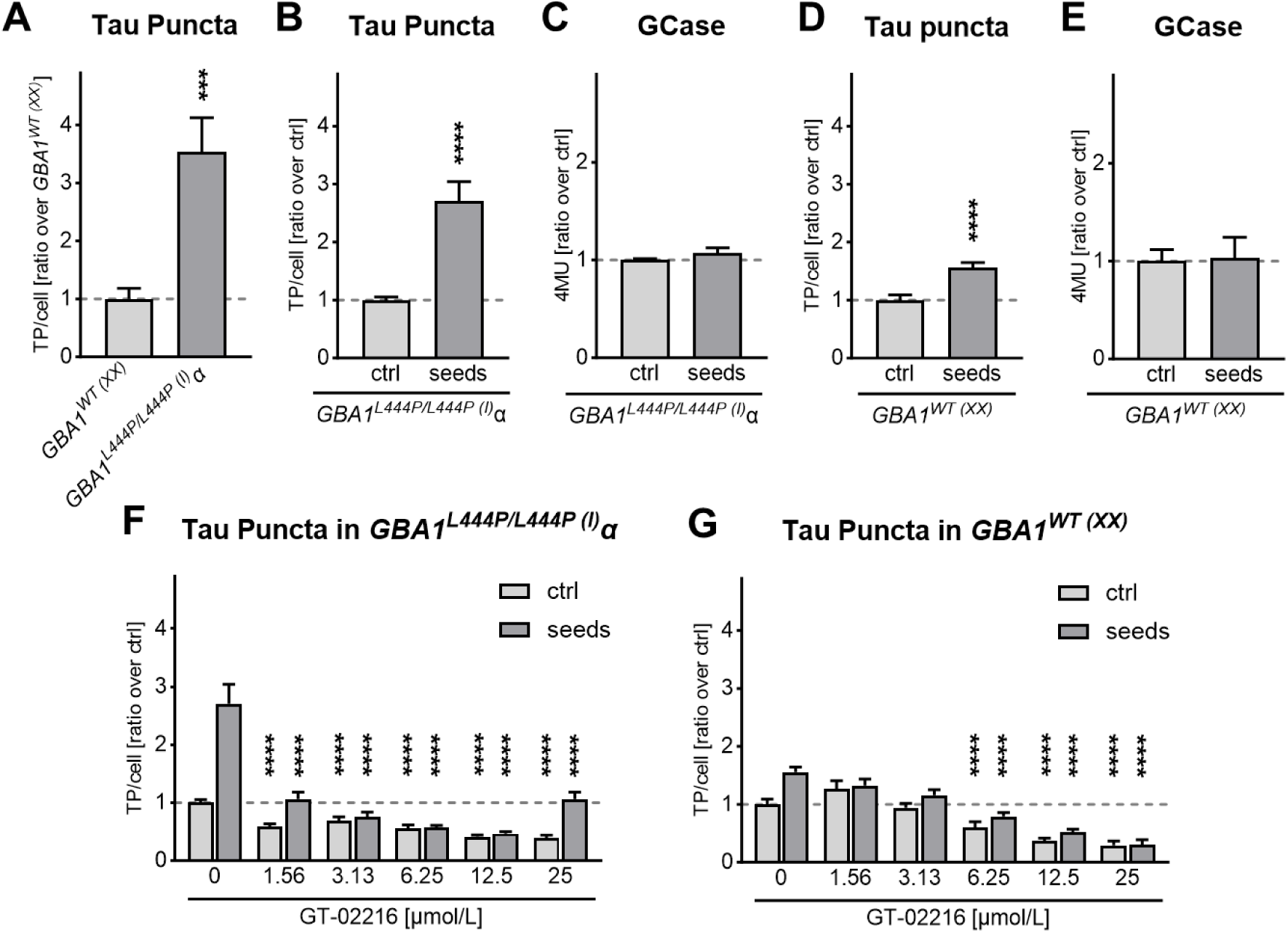
**A.** Quantification of Tau Puncta (TP) in Tau-GBA1^WT(XX)^ and Tau-GBA1^L444P/L444P(I)^α fibroblasts at basal conditions (mean ± sem normalized on Tau-GBA^WT(XX)^, n= 45 fields, 219 cells). Mann-Whitney test, *** p .0004. **B.** Quantification of Tau puncta in Tau-GBA1^L444P/L444P(I)^α fibroblasts treated overnight in the absence (ctrl) or presence (seeds) of Alzheimer’s brain-derived Tau seeds (mean ± sem normalized on ctrl, n= 55-60 fields). Mann-Whitney test, **** p 4×10^-11^. **C.** GCase activity in Tau-GBA1^L444P/L444P(I)^α fibroblasts treated overnight in the absence or presence of Tau seeds (mean ± sem normalized on ctrl, n= 3). Mann-Whitney test, not significant. **D.** as in **B.** for Tau-GBA1^WT(XX)^ fibroblasts (n= 45 fields, 298 cells). Mann-Whitney test, **** p .00001. **E.** as in **C.** for Tau-GBA1^WT(XX)^ fibroblasts (n= 6-9). Mann-Whitney test, not significant. **F.** Dose-dependent reduction of Tau Puncta in Tau-GBA1^L444P/L444P(I)^α fibroblasts or **G.** in GBA1^WT(XX)^ fibroblasts treated for 4 days with the indicated amount of GT-02216 (mean ± sem normalized on vehicle, no seed ctrl, n= 30-45 fields). 2way ANOVA (p <.0001 for GT-02216 concentration and seeds) and Šídák’s multiple tests against vehicle, **** p < .0001.

We then questioned whether Tau accumulation may somehow affect DOs. First, to confirm that Tau-mCherry accumulation occurred in LAMP1-positive DOs in a GT-02216-dependent manner, we determined for each single fibroblast the Pearson’s Correlation Coefficient (PCC) of mCherry biofluorescence with LAMP1 fluorescent staining^35^. Whereas Tau seed-treatment of Tau-*GBA1^L444P/L444P^*^(I)^α fibroblasts significantly increased Tau in LAMP1-positive DOs, the presence of GT-02216 reversed this seed-dependent effect (**Fig. 6A**). Similar, but less pronounced data, were obtained in Tau-*GBA1^WT^*^(XX)^ fibroblasts (**Fig. 6B**). Consistent with our previous findings in normal fibroblasts^24^, Tau seed-treatment increased the number of DOs present in Tau-*GBA1^L444P/L444P^*^(I)^α (**Fig. 6C**) and Tau-*GBA1^WT^*^(XX)^ (**Fig. 6D**) and again, GT-02216 reversed this effect. Possibly, the presence of mutated GCase (compare the light gray bars in **Fig. 6C** and **D**), or seed-induced Tau accumulation in DOs, led to lysosomal malfunction followed by a cellular response in the form of increased DOs biogenesis. On the other hand, the effect of GT-02216 on GCase activity improved lysosomal function, which favored Tau degradation and decreased Tau accumulation in DOs.

**Figure 6.**
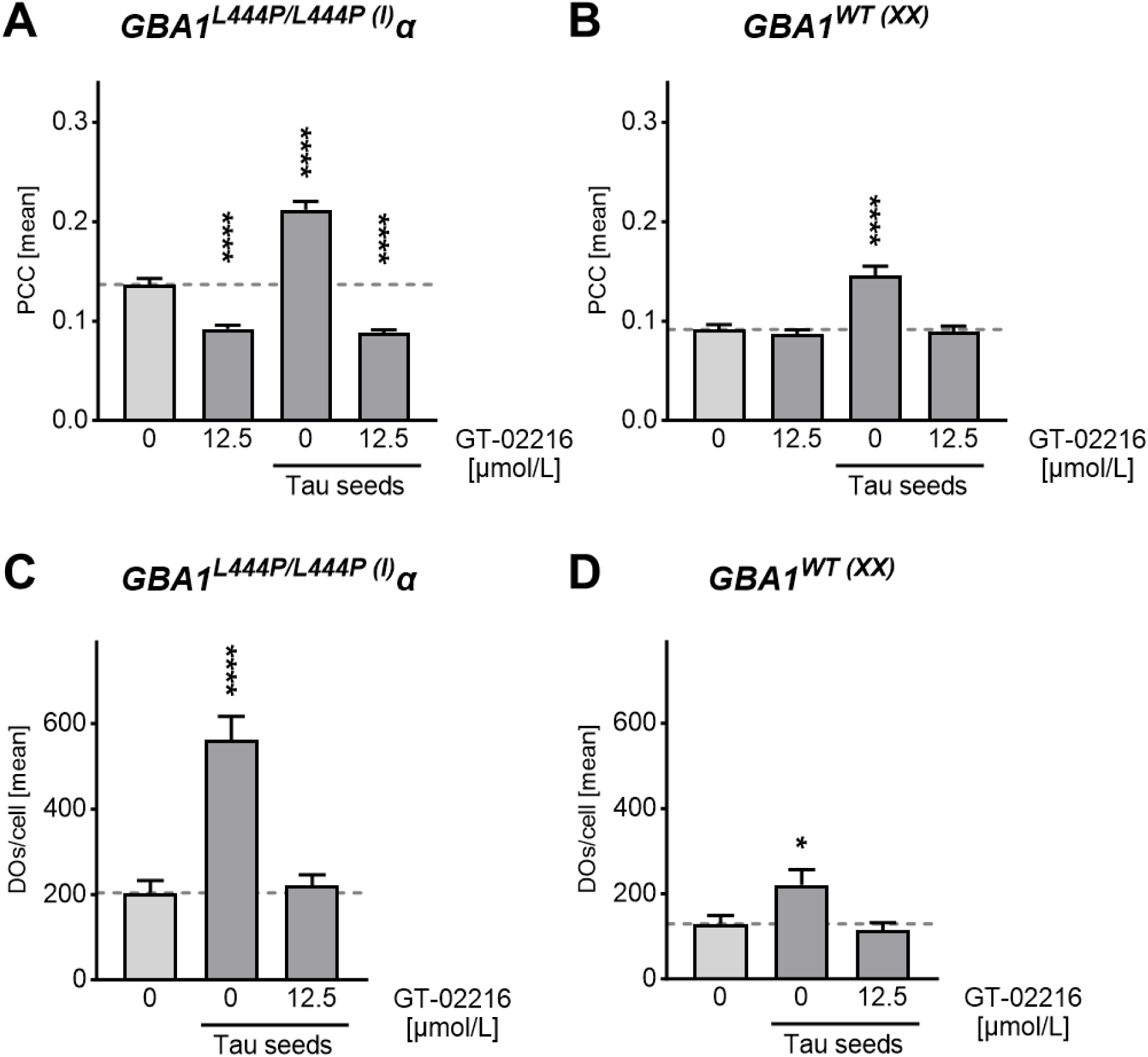
**A.** Pearson’s Correlation Coefficients (PCC) for Tau-mCherry on LAMP1-positive DOs in Tau-GBA1^L444P/L444P(I)^α fibroblasts treated for 4 days with the indicated GT-02216 concentration and overnight in the presence or absence of Tau seeds (mean ± sem, n= 100 cells). **B.** As in **A.** for Tau-GBA1 ^WT(XX)^ fibroblasts. **C.** Number of LAMP1-positive DOs per Tau-GBA1^L444P/L444P(I)^α fibroblast treated as in **A.** (mean ± sem, n= 20 cells). **D.** As in **C.** for Tau-GBA1^WT(XX)^ fibroblasts. Ordinary 1way ANOVA (p <.0001, except for p <.01 in D.) and Dunnett’s multiple tests against the respective controls (light grey bars), * p < .05, **** p < .0001.

### GT-02216 protects rat hippocampal primary neurons challenged with Tau oligomers

To further explore the effects of GT-02216 in a different cellular context, we extended our investigation to a neuronal model. Given that Tau oligomers (TauO) are known to exert neurotoxic effects, we assessed whether GT-02216 could confer neuroprotection in primary rat hippocampal neurons challenged with TauO. Incubation of these cells with TauO at a concentration of 5 μmol/L resulted in a significant decrease in cell viability, consistent with previous findings^36,37^. Recombinant Brain Derived Neurotrophic Factor (BDNF), known to be protective against Tau-related neurotoxicity in several neurodegenerative disorders^38^, was used as a positive control in the assay. BDNF significantly attenuated TauO-induced neuronal death in our model validating the experimental set-up (**Fig. 7A).** GT-02216 treatment also effectively rescued cell viability in the TauO-challenged hippocampal neurons, demonstrating its neuroprotective potential against TauO-induced neurotoxicity (**Fig. 7A**).

**Figure 7.**
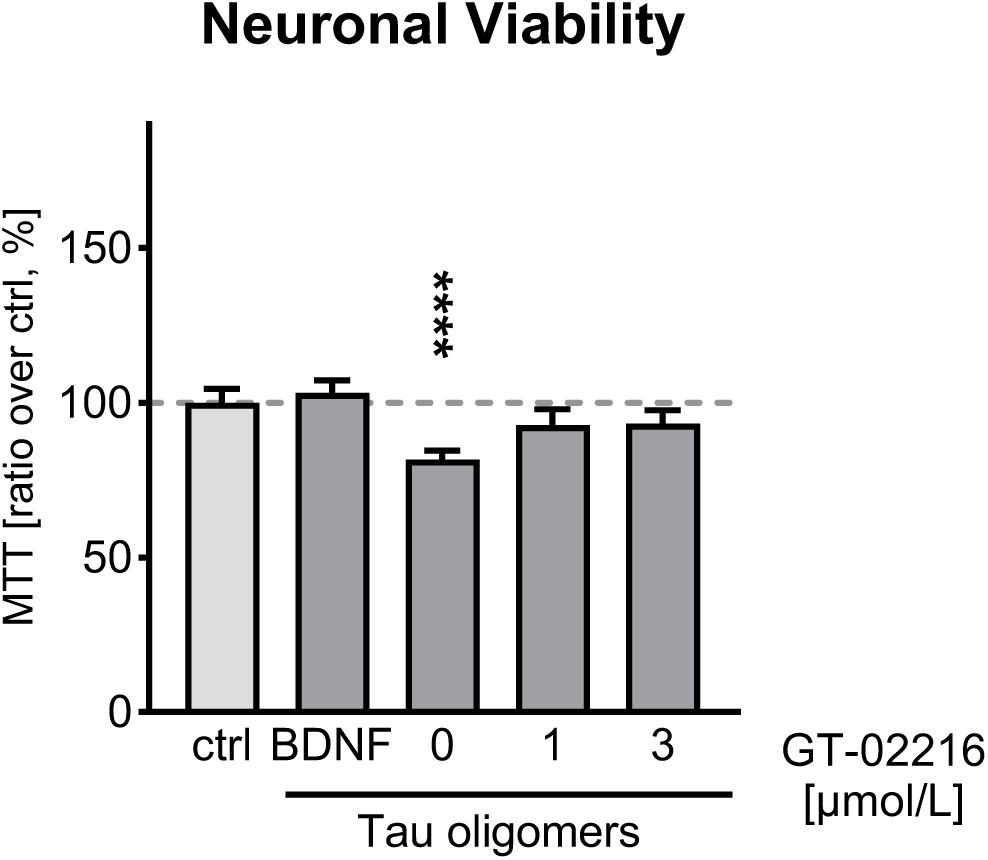
Viability of rat hippocampal primary neurons treated for 2 days with GT-02216 at the indicated concentration or with BDNF before being challenged with 5 µmol/L Tau oligomers for 1d. Cell viability was assessed with the MTT assay (mean ± SD, n= 3-4 wells). Ordinary 1way ANOVA (p <.0001) and Dunnett’s multiple tests against the control, **** p < .0001.

## Discussion

Our study was an initial exploration of the therapeutic potential of the pharmacological GCase-enhancing chaperone GT-02216 in tauopathies. GT-02216 was selected starting from a novel druggable allosteric GCase site identified with the proprietary Gain Therapeutics SEE-Tx® drug-discovery platform^25,27,28^. *In vitro* assays showed GT-02216 binding to GCase that increased GCase activity and reduced endogenous GCase substrates in fibroblasts. The SEE-Tx® computational technology was previously successfully applied to other enzymes linked to LSD^39,40^. Examples are alpha-L-iduronidase deficiency causing mucopolysaccharidosis type I, an inherited lysosomal disease^40^, and mutant β-galactosidase^39,41^ triggering GM1 gangliosidosis and mucopolysaccharidosis type IVB, also known as Morquio B disease^39,41^.

GCase enzymatic dysfunction leads to defects in metabolism and lysosomal stress. We selected several human dermal fibroblasts derived from healthy donors and donors carrying *GBA1* mutations as our model. We first validated the model showing that the activity of GCase was reduced in *GBA1* mutation-carrying fibroblasts when compared to normal fibroblasts, which led to increased amounts of intracellular sphingolipids, the substrates of lysosomal GCase. We then showed that GT-02216 enhanced GCase activity and reduced HexCer accumulation in *GBA1^L444P/L444P^* mutant and wild-type fibroblasts. Fibroblasts carrying the homozygous L444P/L444P mutation, which is associated with severe neuronopathic Gaucher’s disease and increased dementia risk^42^, showed the strongest decrease in GCase activity and were therefore selected for the whole study. To evaluate the pathological accumulation of Tau in this model, we took advantage of a previous study reporting evidence of aberrant Tau accumulation in DOs linked to lysosomal stress in primary human fibroblasts expressing various fluorescent forms of Tau^24^. This adverse process was further propagated by incubating the cells with Alzheimer’s brain-derived Tau seeds as well as in the presence of CBE, a cell-active pharmacological irreversible inhibitor of lysosomal GCase^24^. In agreement with these data, we observed increased seed-induced Tau accumulation in DOs of Gaucher’s fibroblasts expressing defective GCase when compared to wild-type fibroblasts. These data stress the potential implication of GCase deficiency and the accumulation of GCase substrates in abnormal Tau accumulation in DOs. Of relevance in this context, treatment of the cells with GT-02216 efficiently reduced Tau accumulation. Moreover, the rescuing effect was also observed in cells with the wild-type GCase background. We concluded that lysosomal dysfunction associated to GCase deficits in combination with internalized Tau seeds may together contribute to the accumulation of Tau on route to degradation within DOs. However, Tau accumulation and impairment of lysosomal function and lipid metabolism may reciprocally engage in a sequence of harmful events. Improving lysosomal function through pharmacological GCase enhancement may represent a viable strategy to reduce aging-associated cellular stress by indirectly targeting Tau accumulation. The reduction of Tau accumulation in basal and seed-induced conditions after treatment with GT-02216 observed in fibroblasts, acquired further relevance from the finding that GT-02216 rescued the viability of (endogenously Tau expressing) rat hippocampal neurons challenged with exogenous Tau oligomers. This highlighted the critical balance maintained by ALP in neuronal health and showed the potential of GT-02216 as a neuroprotective agent targeting Tau-related neurotoxicity with an effect comparable to that of the neuroprotectant BDNF.

Our study further confirms the emerging crosstalk between ALP dysfunction and neurodegeneration. Mice deficient in autophagy exhibit neuronal accumulation of aggregate-prone proteins and neurodegeneration, demonstrating the crucial role of autophagy in neuronal homeostasis^43,44^. The role of the ALP in neurodegenerative diseases has been extensively studied in recent years. Late-onset neurodegenerative disorders such as Parkinson’s, Huntington’s, and Alzheimer’s diseases are characterized by the accumulation of intracellular aggregates in the brain. Clearance of these aggregates is typically associated with improvement of symptoms^45^, indicating that ALP impairment and accumulation of pathogenic protein forms may together contribute to disease progression. Tau lesions are detected in LSD models such as Gaucher’s disease, Niemann–Pick, Sanfilippo syndrome type B, Christianson syndrome and Fabry’s disease^46–49^. Moreover, restoring mutated GCase activity with the chaperone Ambroxol or through ectopic expression of wild-type GCase delayed Tau and α-synuclein accumulation^50,51^. Thus, correcting ALP defects are appealing therapeutic interventions.

Considering the data showing efficacy of GT-02216 also in wild-type cells, our study prompts additional investigations to further underpin that boosting GCase activity could represent a viable therapeutic strategy to slow-down aging-dependent protein deposition in the general patient population not affected by *GBA1* mutations. Despite the fact that *GBA1* mutations are not linked to Alzheimer’s disease or other neurodegenerative proteinopathies besides Parkinson’s disease^52,53^, altogether, our findings suggest that GT-02216 shows promise as a potential disease-modifying treatment option for aging, Alzheimer’s disease or other tauopathies beyond those directly associated with *GBA1* mutations.

## Materials and Methods

### Binding studies by Surface Plasmon Resonance (SPR)

Human full-length wild-type GCase protein (Cerezyme, Genzyme, Naarden, NL) was immobilized on the SPR CM5 sensor (GE Healthcare, #29149603,) by standard amino coupling using relatively high protein concentration of 100 µg/mL. A nine point 2-fold serial dilution starting from 100 µmol/L GT-02216^26^ (10 mmol/L stock solution in DMSO) was measured by SPR at pH 7.4 in 10 mmol/L HEPES, 5 mmol/L EDTA, 150 mmol/L NaCl and 0.01% Tween-20 or at pH 5.0 in 20 mmol/L Na phosphate, 2.7 mmol/L KCl, 137 mmol/L NaCl, 5 mmol/L tartrate, 0.01% Tween-20. Empty, activated, and deactivated parallel channels on the same SPR sensor were used as reference channels. Raw SPR signals monitored on the active channel with immobilized GCase protein were subtracted with signals monitored on the reference channel (empty sensor surface) and further subtracted with the signal monitored for the running buffer (double referenced), and finally corrected for DMSO signal mismatch between sample and running buffer. To extract binding affinity values, the plotted SPR data were further fitted with the four-parameter logarithmic dose-response equation without constraint (GraphPad Prism version 10.2.3).

### 4MU GCase activity assay

Fibroblasts were seeded at 5×10^3^ cells/well on poly-D-lysine (Sigma-Aldrich, #P6407) coated 96-multiwell plates (ThermoFisher Scientific™, #167008). For 4MU determination, the cell culture medium was removed, cells were washed with PBS and then supplemented with 5 mmol/L 4-methylumbelliferyl-beta-D-glucopyranoside (Apollo Scientific, #BIM1097) in 0.1 mol/L acetate buffer pH 4. The reaction was stopped with 100 mmol/L Na glycine pH 10.7 after 1 h at 37°C. 4MU fluorescence was determined at ex/em 340/460 nm with a fluorescence plates reader (TECAN, Infinite 200 Pro^®^).

### Human dermal fibroblasts

Human primary *GBA1^WT^*^(XX)^ fibroblasts were isolated from a dermal biopsy obtained from a healthy 30-year-old female^24^. *GBA1^L444P/L444P^*^(I)^α Gaucher type I fibroblasts (Coriell, #GM10915) were originally isolated from a 7-year-old male and are homozygote for a T>C transition at nucleotide 1448 in exon 10 of the *GBA1* gene. Additional human fibroblasts used in this study were *GBA1^WT^*^(XY)^ male (Coriell, #GM03377), *GBA1^N370S/V394L^*^(I)^ Gaucher type I (Coriell, #GM01607), *GBA1^N370S/84GG^*^(I)^ Gaucher type I (Coriell, #GM00372), *GBA1^N188S/S107L^*^(III)^ Gaucher type III (Telethon Biobank, #20843), *GBA1^L444P/F213I^*^(III)^ Gaucher type III (Telethon Biobank, #21142), *GBA1^L444P/R496C^*^(III)^ Gaucher type III (Telethon Biobank, #20624), *GBA1^L444P/L444P^*^(III)^ Gaucher type III (Telethon Biobank, #20526), *GBA1^L444P/L444P^*^(II)^ Gaucher type II (Coriell, #GM08760), and *GBA1^L444P/L444P^*^(I)^β Gaucher type I (Coriell, #GM07968).

Fibroblasts were cultured in an incubator at 37°C, 5% CO_2_ in DMEM/GlutaMAX^TM^ (Gibco, #61965-026) containing 15% FBS (Sigma-Aldrich, #0001668922), 1% Pen/Strep (Gibco, #15140-122), and MEM NEAA (Gibco, #1140-035). Fibroblasts were transduced for doxycycline-inducible Tau-mCherry as described^24^.

### Fibroblast treatments

Tau expression was routinely induced for 4 days with 0.3 µg/mL doxycycline (Sigma-Aldrich, #D9891). GT-02216 was incubated with cells for 4 days. Independently of the compound concentration, the vehicle DMSO was kept constant. Tau seeds were enriched from frozen Alzheimer’s disease frontal cortex samples as described^24^. The tissues were obtained from The Netherlands Brain Bank, Netherlands Institute for Neuroscience, Amsterdam (www.brainbank.nl). Anonymized donors signed a written informed consent for brain autopsy and further use of tissue and clinical information for research purposes. Tau seeds were directly supplemented to the cell culture media for 1 day before the cells were gently washed with PBS (Gibco, #10010023), cell nuclei were stained with 2.5 μg/mL Hoechst 33342 (Invitrogen, #H3570) for 10 min, 37°C, followed by gentle washes in complete medium and PBS, before fixation for further analysis.

For Tau accumulation analysis, fibroblasts seeded at 30×10^3^ cells/well on poly-D-lysine (Sigma-Aldrich, #P6407) coated 8-well microscope slides (Ibidi, #80826-IBI) were fixed in 2% formaldehyde in PBS for 15 min at room temperature, washed three times with 500 mmol/L glycine in PBS and then three times with PBS.

For LAMP1 immunostaining analysis, fibroblasts were fixed in 4% formaldehyde in PBS for 5 min at 37°C followed by cold 2% formaldehyde in 6.25% methanol (Sigma-Aldrich, #32213) in PBS for 1 min at room temperature. Cells were washed three times with 500 mmol/L glycine in PBS and two times in PBS. Unspecific binding was blocked with 5% NGS (Sigma-Aldrich, #N2513) in PBS for 1 h at room temperature and three washes in PBS. Staining with 0.4 µg/ml LAMP1 primary antibody (SantaCruz Biotechnology, Inc., #SC20011) was done in 0.5% NGS in PBS for 1 h at room temperature, followed by three washes in PBS and the addition of 2 µg/ml AlexaFluor^TM^ 488 (Invitrogen, #A11001) secondary antibody and 1 µg/ml DAPI (Sigma-Aldrich, #D9542) for 1 h at room temperature. After the last incubation, cells were washed three times and stored in PBS.

Fluorescence images were acquired with a laser confocal microscope (Leica Microsystems, STELLARIS 8), using specific laser detectors for Hoechst 33342 (ex/em 352/455 nm), DAPI (ex/em: 360/460 nm), AlexaFluor^TM^ 488 (ex/em: 499/520 nm) and mCherry (ex/em 587/610 nm).

Image analysis and processing of the raw data were performed with Fiji/ImageJ v1.54 software. Pearson’s Correlation Coefficient (PCC) was determined with ImageJ JACoP v2.0 plugin for dual-color colocalization of Tau and LAMP1. Acquisition settings were kept constant across all conditions analyzed and acquired images were processed with the exact same parameters, including background subtraction. Quantification of the percentage of cells with a Tau puncta phenotype was normalized with the total number of DAPI positive nuclei. Single Tau puncta were manually identified and analyzed with the ImageJ software^24,35^.

### Hexosylceramide Assay

Primary human *GBA1^WT (XX)^* fibroblasts were seeded at 2.5×10^5^ cells/T25 flasks (Falcon^®^, #353808) in DMEM+GlutaMAX^TM^ containing 15% FBS and 1% PenStrep and incubated at 37°C, 5% CO_2_ for 1 day. Cell media were then replaced with fresh medium containing the desired concentration of GT-02216. After 10 days, cells were detached, collected into two 2 mL tubes for each flask, and placed in ice. After centrifugation at 13000 rpm at 4°C for 5 min, the cell pellets were rinsed with 1 mL cold PBS and collected into a single tube for each sample. Samples were centrifuged at 13000 rpm at 4°C for 2 min and cells pellets stored at -80°C until further analysis.

Cell pellets were dissolved in 150 μL 0.5% formic acid in methanol supplemented with 1 µmol/L C18 galactosylceramide-d_35_ (d18:1/18:0-d_35_) (GalCer-d_35_; Cayman, #24467) as internal standard. The samples were shaken for 1 h, 1x sonicated (4 sec, 30% amplitude) and centrifuged at 13000 g at 4°C for 5 min. Sphingolipid analysis of supernatants was performed by UPLC-MS. An InfinityLab Poroshell 120 HILIC, 2.1×100 mm,1.9 μm column was used at a flow rate of 0.5 mL/min with solvent A composed of 5 mM ammonium formate, 0.1% formic acid in water and solvent B composed of 5 mM ammonium formate, 0.1% formic acid in acetonitrile/water/methanol 95:2.5:2.5. The gradient was programmed for 0-0.1 min 95% B; 0.1-1.0 min 95–0% B, 1.0-2.0 min 0% B, 2.0-2.1 min 0-95% B, and 2.1–2.5 min 95% B. The mass spectrometer (Agilent 1290 Infinity II / QQQ G6475) was operated in a positive ESI mode with a capillary voltage of 4600 V, a nozzle voltage of 1700 V, a gas temperature of 280°C, a gas flow of 10.5 L/min and a nebulizer at 33 psi. The sheath gas temperature and gas flow were at 270°C and at 11.5 L/min, respectively. Hexosylceramide (HexCer) was analyzed in multiple reaction monitoring mode; 728.6 > 710.5 m/z transition was quantified.

### Rat Primary Hippocampal Neurons

Hippocampal neurons were prepared as described^54–56^(ETAP–Lab; https://www.etap-lab.com; France). Briefly, the rat embryos were delicately extracted, brains were quickly collected, and the hippocampus dissected from each brain using a binocular magnifier. Single cells were obtained after dissociating the hippocampal tissues by enzymatic digestion and mechanical separation. Dissociated neurons were plated at 80 x10^3^ cells/well in 48-well plates pre-coated with 15 µg/mL poly-ornithine (Sigma-Aldrich, #P4957) and 1 µg/mL laminin (Sigma-Aldrich, #11243217001). Cells were cultured at 37°C in a humidified 5% CO_2_ atmosphere in serum-free neurobasal medium (Invitrogen, #21103049) and 2% B27 supplement (Invitrogen, #17504044). Starting from day 4, neurons were preincubated with GT-02216, vehicle (diluted DMSO) or brain derived neurotrophic factor (BDNF; Sigma-Aldrich, #B3795, lot #0000100311) at 37°C for 48 h. On day 6, Tau oligomers were added into the cell media for 24 h. The Tau oligomers were prepared according to the ETAP-Lab’s protocol starting from recombinant human Tau_2N4R_ monomers. Tau oligomers contained a mixture of trimers, low molecular weight oligomers and remaining monomeric forms of the protein.

Primary neuron viability was evaluated at ETAP-Lab (https://www.etap-lab.com; France) by the MTT assay (3-(4,5-dimethylthiazol-2-yl)-2,5-diphenyltetrazolium bromide) at 24 h post the addition of Tau oligomers. Briefly, cells were incubated at 37°C for 1 h with 0.5 mg/mL MTT; medium was removed, and cells lysed in 100% DMSO. After complete solubilization, formazan absorbance was recorded at 500 nm with a spectrophotometer (Molecular Devices, SpectraMax® i3x).

### Statistics and reproducibility

Statistical analysis was performed with GraphPad Prism version 10.2.3 with at least three independent biological replicates. Most quantifications are reported as fold over control/untreated conditions, unless otherwise indicated in the graphs.

## Acknowledgments

We thank the whole laboratory for support and advice during this study. We thank the Microscopy facility of the Bellinzona Institutes of Science (BIOS+) for support during the executions and analysis of the experiments. We thank Elena Cubero, ETAP LAB, USC and PCB. Research was mainly supported by the Innovation Project 35449.1 IP-LS from the Swiss Innovation Agency. The Paganetti’s lab is founded by the Gelu Foundation, the Mecri Foundation and The Charitable Gabriele Foundation.

## Author contributions

Conceptualization: PP, SP, MM, MB, NP-C, AMG-C

Methods and investigations: MC, EP, IF, SP, AR, AD, BC-FG

Supervision: PP, SP, NP-C, JT, AMG-C

Writing: PP, SP, MC, SC-C (original draft), all co-authors (review & editing)

## Data availability

all raw data can be found in the supplementary files.

## References

1. Chiti, F. & Dobson, C. M. Protein Misfolding, Amyloid Formation, and Human Disease: A Summary of Progress Over the Last Decade. Annu. Rev. Biochem. 86, 27–68 (2017).

2. Goedert, M. & Spillantini, M. G. A century of Alzheimer’s disease. Science 314, 777–781 (2006).

3. Nisbet, R. M. & Götz, J. Amyloid-β and Tau in Alzheimer’s Disease: Novel Pathomechanisms and Non-Pharmacological Treatment Strategies. J. Alzheimers Dis. JAD 64, S517–S527 (2018).

4. Spillantini, M. G. & Goedert, M. Neurodegeneration and the ordered assembly of α-synuclein. Cell Tissue Res. 373, 137–148 (2018).

5. Giovedì, S., Ravanelli, M. M., Parisi, B., Bettegazzi, B. & Guarnieri, F. C. Dysfunctional Autophagy and Endolysosomal System in Neurodegenerative Diseases: Relevance and Therapeutic Options. Front. Cell. Neurosci. 14, 602116 (2020).

6. Finkbeiner, S. The Autophagy Lysosomal Pathway and Neurodegeneration. Cold Spring Harb. Perspect. Biol. 12, a033993 (2020).

7. Nixon, R. A. The role of autophagy in neurodegenerative disease. Nat. Med. 19, 983–997 (2013).

8. Fraldi, A., Klein, A. D., Medina, D. L. & Settembre, C. Brain Disorders Due to Lysosomal Dysfunction. Annu. Rev. Neurosci. 39, 277–295 (2016).

9. Monaco, A. & Fraldi, A. Protein Aggregation and Dysfunction of Autophagy-Lysosomal Pathway: A Vicious Cycle in Lysosomal Storage Diseases. Front. Mol. Neurosci. 13, 37 (2020).

10. Shachar, T. et al. Lysosomal storage disorders and Parkinson’s disease: Gaucher disease and beyond. Mov. Disord. Off. J. Mov. Disord. Soc. 26, 1593–1604 (2011).

11. Robak, L. A. et al. Excessive burden of lysosomal storage disorder gene variants in Parkinson’s disease. Brain J. Neurol. 140, 3191–3203 (2017).

12. Behl, T. et al. Cross-talks among GBA mutations, glucocerebrosidase, and α-synuclein in GBA-associated Parkinson’s disease and their targeted therapeutic approaches: a comprehensive review. Transl. Neurodegener. 10, 4 (2021).

13. McNeill, A. et al. Ambroxol improves lysosomal biochemistry in glucocerebrosidase mutation-linked Parkinson disease cells. Brain J. Neurol. 137, 1481–1495 (2014).

14. Gegg, M. E. et al. Glucocerebrosidase deficiency in substantia nigra of parkinson disease brains. Ann. Neurol. 72, 455–463 (2012).

15. Grigor’eva, E. V. et al. Biochemical Characteristics of iPSC-Derived Dopaminergic Neurons from N370S GBA Variant Carriers with and without Parkinson’s Disease. Int. J. Mol. Sci. 24, 4437 (2023).

16. Emelyanov, A. et al. Increased α-Synuclein Level in CD45+ Blood Cells in Asymptomatic Carriers of GBA Mutations. Mov. Disord. Off. J. Mov. Disord. Soc. 36, 1997–1998 (2021).

17. Kopytova, A. E. et al. Could Blood Hexosylsphingosine Be a Marker for Parkinson’s Disease Linked with GBA1 Mutations? Mov. Disord. Off. J. Mov. Disord. Soc. 37, 1779– 1781 (2022).

18. Murphy, K. E. et al. Reduced glucocerebrosidase is associated with increased α-synuclein in sporadic Parkinson’s disease. Brain J. Neurol. 137, 834–848 (2014).

19. Thomas, R., Moloney, E. B., Macbain, Z. K., Hallett, P. J. & Isacson, O. Fibroblasts from idiopathic Parkinson’s disease exhibit deficiency of lysosomal glucocerebrosidase activity associated with reduced levels of the trafficking receptor LIMP2. Mol. Brain 14, 16 (2021).

20. Du, T.-T. et al. GBA deficiency promotes SNCA/α-synuclein accumulation through autophagic inhibition by inactivated PPP2A. Autophagy 11, 1803–1820 (2015).

21. Mazzulli, J. R. et al. Gaucher disease glucocerebrosidase and α-synuclein form a bidirectional pathogenic loop in synucleinopathies. Cell 146, 37–52 (2011).

22. Yap, T. L. et al. Alpha-synuclein interacts with Glucocerebrosidase providing a molecular link between Parkinson and Gaucher diseases. J. Biol. Chem. 286, 28080–28088 (2011).

23. Muñoz, S. S., Petersen, D., Marlet, F. R., Kücükköse, E. & Galvagnion, C. The interplay between Glucocerebrosidase, α-synuclein and lipids in human models of Parkinson’s disease. Biophys. Chem. 273, 106534 (2021).

24. Piovesana, E. et al. Tau accumulation in degradative organelles is associated to lysosomal stress. Sci. Rep. 13, 18024 (2023).

25. Alonso, X. B., Garcia, D. A. & Schmidtke, P. Method of binding site and binding energy determination by mixed explicit solvent simulations. Patent WO2013092922A2 (2012).

26. Collazo, A. M. G., Jorda, E. C., Bellotto, M. & Iglesias, E. F. Heteroaryl compounds and therapeutic uses thereof in conditions associated with the alteration of the activity of beta-glucocerebrosidase. Patent WO2021105908A1 (2021).

27. Alvarez-Garcia, D. & Barril, X. Molecular simulations with solvent competition quantify water displaceability and provide accurate interaction maps of protein binding sites. J. Med. Chem. 57, 8530–8539 (2014).

28. Alvarez-Garcia, D., Schmidtke, P., Cubero, E. & Barril, X. Extracting Atomic Contributions to Binding Free Energy Using Molecular Dynamics Simulations with Mixed Solvents (MDmix). Curr. Drug Discov. Technol. 19, 62–68 (2022).

29. Shaaltiel, Y. et al. Production of glucocerebrosidase with terminal mannose glycans for enzyme replacement therapy of Gaucher’s disease using a plant cell system. Plant Biotechnol. J. 5, 579–590 (2007).

30. Ruiz-Carmona, S. et al. rDock: a fast, versatile and open source program for docking ligands to proteins and nucleic acids. PLoS Comput. Biol. 10, e1003571 (2014).

31. Schuck, P. Use of surface plasmon resonance to probe the equilibrium and dynamic aspects of interactions between biological macromolecules. Annu. Rev. Biophys. Biomol. Struct. 26, 541–566 (1997).

32. Patching, S. G. Surface plasmon resonance spectroscopy for characterisation of membrane protein-ligand interactions and its potential for drug discovery. Biochim. Biophys. Acta 1838, 43–55 (2014).

33. Premkumar, L. et al. X-ray structure of human acid-beta-glucosidase covalently bound to conduritol-B-epoxide. Implications for Gaucher disease. J. Biol. Chem. 280, 23815–23819 (2005).

34. Rocha, E. M. et al. Sustained Systemic Glucocerebrosidase Inhibition Induces Brain α-Synuclein Aggregation, Microglia and Complement C1q Activation in Mice. Antioxid. Redox Signal. 23, 550–564 (2015).

35. Pedrioli, G. et al. Tau Seeds in Extracellular Vesicles Induce Tau Accumulation in Degradative Organelles of Cells. DNA Cell Biol. 40, 1185–1199 (2021).

36. Flach, K. et al. Tau oligomers impair artificial membrane integrity and cellular viability. J. Biol. Chem. 287, 43223–43233 (2012).

37. Usenovic, M. et al. Internalized Tau Oligomers Cause Neurodegeneration by Inducing Accumulation of Pathogenic Tau in Human Neurons Derived from Induced Pluripotent Stem Cells. J. Neurosci. Off. J. Soc. Neurosci. 35, 14234–14250 (2015).

38. Jiao, S.-S. et al. Brain-derived neurotrophic factor protects against tau-related neurodegeneration of Alzheimer’s disease. Transl. Psychiatry 6, e907 (2016).

39. Rudinskiy, M. et al. Validation of a highly sensitive HaloTag-based assay to evaluate the potency of a novel class of allosteric β-Galactosidase correctors. PloS One 18, e0294437 (2023).

40. Cubero, E. et al. Discovery of allosteric regulators with clinical potential to stabilize alpha-L-iduronidase in mucopolysaccharidosis type I. PloS One 19, e0303789 (2024).

41. Condori, J. et al. Enzyme replacement for GM1-gangliosidosis: Uptake, lysosomal activation, and cellular disease correction using a novel β-galactosidase:RTB lectin fusion. Mol. Genet. Metab. 117, 199–209 (2016).

42. Mahoney-Crane, C. L. et al. Neuronopathic GBA1L444P Mutation Accelerates Glucosylsphingosine Levels and Formation of Hippocampal Alpha-Synuclein Inclusions. J. Neurosci. Off. J. Soc. Neurosci. 43, 501–521 (2023).

43. Komatsu, M. et al. Loss of autophagy in the central nervous system causes neurodegeneration in mice. Nature 441, 880–884 (2006).

44. Hara, T. et al. Suppression of basal autophagy in neural cells causes neurodegenerative disease in mice. Nature 441, 885–889 (2006).

45. Yamamoto, A. & Simonsen, A. The elimination of accumulated and aggregated proteins: a role for aggrephagy in neurodegeneration. Neurobiol. Dis. 43, 17–28 (2011).

46. Sardi, S. P. et al. Augmenting CNS glucocerebrosidase activity as a therapeutic strategy for parkinsonism and other Gaucher-related synucleinopathies. Proc. Natl. Acad. Sci. U. S. A. 110, 3537–3542 (2013).

47. Pacheco, C. D., Elrick, M. J. & Lieberman, A. P. Tau normal function influences Niemann-Pick type C disease pathogenesis in mice and modulates autophagy in NPC1-deficient cells. Autophagy 5, 548–550 (2009).

48. Ohmi, K. et al. Sanfilippo syndrome type B, a lysosomal storage disease, is also a tauopathy. Proc. Natl. Acad. Sci. U. S. A. 106, 8332–8337 (2009).

49. Clarke, J., Kayatekin, C., Viel, C., Shihabuddin, L. & Sardi, S. P. Murine Models of Lysosomal Storage Diseases Exhibit Differences in Brain Protein Aggregation and Neuroinflammation. Biomedicines 9, 446 (2021).

50. Yang, S. Y., Taanman, J.-W., Gegg, M. & Schapira, A. H. V. Ambroxol reverses tau and α-synuclein accumulation in a cholinergic N370S GBA1 mutation model. Hum. Mol. Genet. 31, 2396–2405 (2022).

51. Bae, E.-J. et al. Glucocerebrosidase depletion enhances cell-to-cell transmission of α-synuclein. Nat. Commun. 5, 4755 (2014).

52. Tsuang, D. et al. GBA mutations increase risk for Lewy body disease with and without Alzheimer disease pathology. Neurology 79, 1944–1950 (2012).

53. Wen, Y.-F. et al. Mutations in GBA, SNCA, and VPS35 are not associated with Alzheimer’s disease in a Chinese population: a case-control study. Neural Regen. Res. 17, 682–689 (2022).

54. Garcia, P. et al. Ciliary neurotrophic factor cell-based delivery prevents synaptic impairment and improves memory in mouse models of Alzheimer’s disease. J. Neurosci. Off. J. Soc. Neurosci. 30, 7516–7527 (2010).

55. Roby, M. H. et al. Enzymatic production of bioactive docosahexaenoic acid phenolic ester. Food Chem. 171, 397–404 (2015).

56. Colin, J. et al. Improved Neuroprotection Provided by Drug Combination in Neurons Exposed to Cell-Derived Soluble Amyloid-β Peptide. J. Alzheimers Dis. JAD 52, 975–987 (2016).

